# Developmentally programmed loss of long-range Polycomb interactions is regulated by cohesin

**DOI:** 10.64898/2026.09.15.749901

**Authors:** Valeriia Smialkovska, Ann-Kristin Dicke, Angelika Feldmann

## Abstract

Distal regulatory elements (DREs), such as enhancers, can regulate genes across megabase-long distances, presumably via coming into close spatial proximity. The establishment of new transcriptional programmes during cell type transitions is associated with widespread rewiring of the spatial organisation of the genome, including gain and loss of chromatin interactions. Extensive effort has been invested into understanding how chromatin interactions are formed during development, yet the mechanisms underlying their developmental loss remain largely unclear. By leveraging chromatin accessibility-assisted footprinting, acute protein degradation and chromatin conformation capture, we show that loss of promoter interactions cannot be explained by reduced binding of sequence-specific transcription factors (TFs). Instead, we identify a subset of interactions that depend on cohesin for programmed developmental disruption. These sites are characterized by high Polycomb enrichment and TF occupancy and engage in strong long-range interactions that undergo extensive differentiation-dependent rewiring. Preventing interaction loss by acute cohesin degradation results in the preferential downregulation of associated genes. Together, these results suggest that cohesin indirectly regulates developmental loss of Polycomb interactions by enabling the acquisition of other potentially regulatory contacts in a process that may shape transcriptional programs during cell type transitions.

## Introduction

The establishment and maintenance of unique cell type identity relies on the activity of distinct sets of genes, whose expression is coordinated by distal regulatory elements (DREs), including enhancers^1^. DREs can influence their target gene promoters from genomic distances of up to several megabases^2–4^, a process thought to occur upon coming into close spatial proximity^5^.

Many DNA binding factors have been proposed to regulate these physical interactions. A prominent example is the interplay between cohesin and CTCF, where cohesin acts as a loop extruder, reeling in DNA until it is halted and stabilised by CTCF bound at convergently oriented sites^6–11^. In ensemble Hi-C maps, this process is visible as topologically associated domains (TADs), appearing as regions of increased contact frequency relative to their surroundings^12,13^. Interestingly, cohesin and CTCF are frequently enriched at active promoters and enhancers^14^. In support of a functional explanation, ongoing transcription^15–18^ as well as enhancer activation^19,20^ have been proposed to act as extrusion boundaries and sites of cohesin recruitment, respectively. Current consensus suggests that cohesin-mediated loop extrusion creates opportunities but is not the sole determinant for regulatory encounters between promoters and enhancers^21^. Alongside its role in inducing genomic proximity, cohesin has also been implicated in constant disruption of very long-range interactions between super-enhancers, Polycomb sites and A/B compartments^7,22,23^.

Other factors involved in shaping chromatin contacts include chromatin-modifying complexes, such as the Polycomb Repressive Complexes PRC1^24^ and PRC2^25^, transcriptional machinery^15–18^ and co-transcription complexes including several forms of the Mediator complex^26,27^. Surprisingly, only a small number of transcription (co-)factors have been shown to affect physical proximity. These factors include Ldb1^28,29^, Pax5^30^, Foxp3^31^, Oct4 and Nanog^32^, Klf4^33^ and YY1^34^, although the degree of evidence for their active and direct effect on looping varies between proteins and studies^21^.

In agreement with a regulatory role of 3D genome structure, chromatin undergoes extensive rewiring during the establishment of new transcriptional programs, as observed in the course of cell differentiation^35–38^, embryonic development in Drosophila^39^ and mice^40^, or activity-related induction^2,41^. Rewiring during cell type transitions is characterized by stable and transient loss and gain of interactions between gene promoters and DREs. While interactions that are gained by genes that become induced during differentiation are characterized by the activity-associated histone modifications H3K27ac and/or H3K4me1^35,38–40^, loss of interactions across these genes involves sites with high H3K27me3 levels^38,40^, raising the possibility that they are repressive.

Interestingly, rewiring of induced genes during retinoic acid mediated differentiation of embryonic stem cells (ESCs) is dominated by interaction loss^38^, whereby the associated DREs are required for efficient gene activation, in some cases even despite bearing repressive chromatin signatures^26^. This suggests that interaction loss may play an important role in the establishment of transcriptional programs. Whether this developmental loss of promoter-DRE contacts is indeed required for correct gene expression remains unclear. Studying its impact in detail is currently limited, largely because the relevant regulatory factors have not been identified yet.

Here, we use a combination of ATAC footprinting, acute protein degradation and chromatin conformation capture to determine the mechanisms underlying developmental loss of interactions. We show that loss of promoter interactions does not coincide with reduced binding of sequence-specific TFs. Instead, we find that it depends on the presence of cohesin for a subset of sites with high Polycomb enrichment and strong long-range interactions, suggesting an active, developmentally programmed process. Such interactions are further characterized by increased differentiation-dependent rewiring, which correlates with a high TF occupancy. Moreover, impaired loss of cohesin-dependent interactions coincides with reduced transcription of the associated genes. These results are compatible with the possibility that cohesin indirectly regulates loss of repressive interactions by enabling the gain of other regulatory interactions to shape transcription upon differentiation.

## Results

### Developmental loss of interactions occurs independently of a reduction in TF binding

To unbiasedly detect TFs potentially responsible for the developmental loss of interactions, we have chosen a retinoic acid (RA)-dependent differentiation of mouse ESCs in which we have previously measured interaction dynamics at scale^38^. This time-course is characterized by a rapid loss of interactions, with the majority of interaction loss occurring within the first two hours of RA treatment (**Fig. 1A**). To ensure an efficient and complete capturing of interactions lost during early development, we decided to focus our analysis on an early time point (4h).

**Figure 1.**
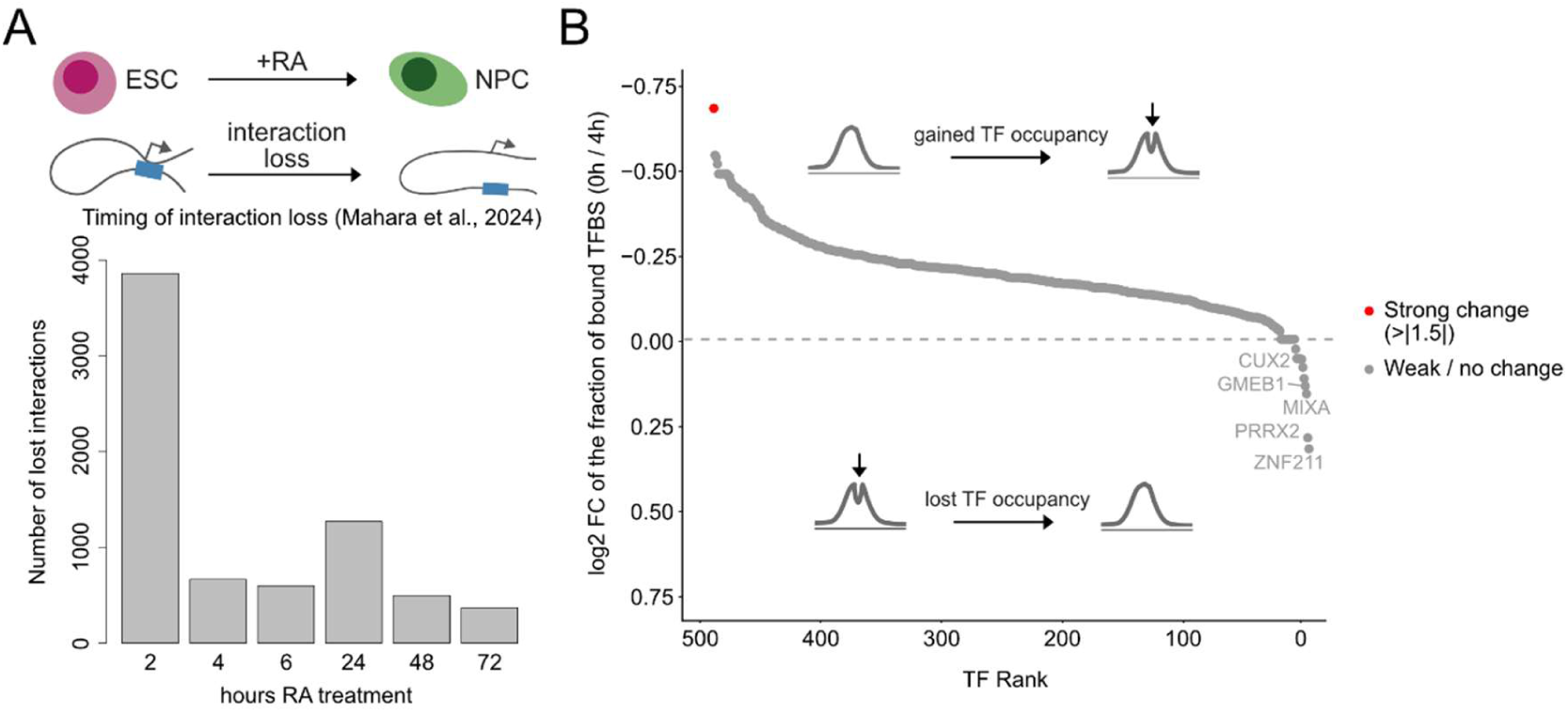
Developmental loss of interactions occurs independently of a reduction in TF binding. **(A)** Barplot showing that developmental loss of interactions mainly occurs at early time-points after RA-induced ESC differentiation^38^. **(B)** Comparison of the fraction of bound transcription factor binding sites (TFBSs) within anchors of developmentally lost interactions as identified by ATAC-seq footprinting. TFs are ranked by differential enrichment between 0h and 4h RA treatment.

During this early differentiation, we hypothesized that the simplest mechanism for developmental interaction loss is a reduction in the binding of specific looping factors at interaction anchors. Aiming to test this hypothesis in a global and unbiased manner, we conducted ATAC-seq before and after four hours of differentiation. We detected a total of 227,960 high-consensus ATAC peaks, of which 1,697 were up-and 4,432 downregulated (DESeq2 padj < 0.05 and fold change ≥ 1.5-fold) after the differentiation (**Fig. S1A**). 98.9% (1,663/1,682) of ATAC peaks that overlapped developmentally lost interaction anchors remained stable (**Fig. S1A-C**), suggesting that interaction loss was not accompanied by a general reduction in accessibility. To identify if interaction loss is linked to the differential binding of TFs, we performed accessibility-assisted footprinting analysis. While we identified one TF (TP53) whose binding increased at sites with lost interactions during the differentiation (**Fig. 1B**, red dot), we could not detect any significantly reduced TF footprints at a cut-off of 1.5-fold (**Fig. 1B**). These results suggest that the developmental loss of interactions is not simply a consequence of reduced binding of sequence-specific TFs, but instead may be regulated by a more general mechanism.

### Cohesin is required for the efficient loss of a subset of promoter interactions during embryonic stem cell differentiation

In search of a mechanism for developmental loop disruption, we aimed to identify DNA-bound factors that have been previously found to be involved in such activity in steady-state cells. Although two DNA-binding factors (cohesin and CTCF) have been previously implicated in prevention or disruption of long-range interactions^7,9,23,42,43^, cohesin is the only complex proposed to actively disrupt interactions in ESCs, as well as short-lived transient interactions during differentiation^38^. We therefore hypothesized that cohesin may also play a role in disrupting pre-existing interactions in a dynamic system, such as during the RA-induced differentiation.

To test whether loss of promoter interactions during differentiation requires cohesin, we reanalyzed our previously published Capture-C datasets^38^, for which we acutely degraded cohesin followed by a four-hour RA treatment (**Fig. 2A**). These Capture-C datasets target a total of 208 gene promoters, including genes that become induced during the differentiation, pluripotent control genes and genes that remain stably expressed throughout the differentiation. Consistent with our previous data^38^, we observed more loss (n=586) than gain of interactions (n=342) in wild-type cells (**Fig. 2B** upper panel). Upon RA-treatment, 91% of these interactions weakened in cohesin-depleted cells compared to cells with normal cohesin levels, as expected given its loop-promoting function (**Fig. 2B** lower panel and **S2A-C**). However, a subset of interactions appeared significantly stronger following cohesin degradation (n=80, 1%), which highly overlapped with developmentally lost interactions (**Fig. 2B**, lower panel and **S2B**). These interactions depended on cohesin for disruption, suggesting an active, developmentally programmed mechanism.

**Figure 2.**
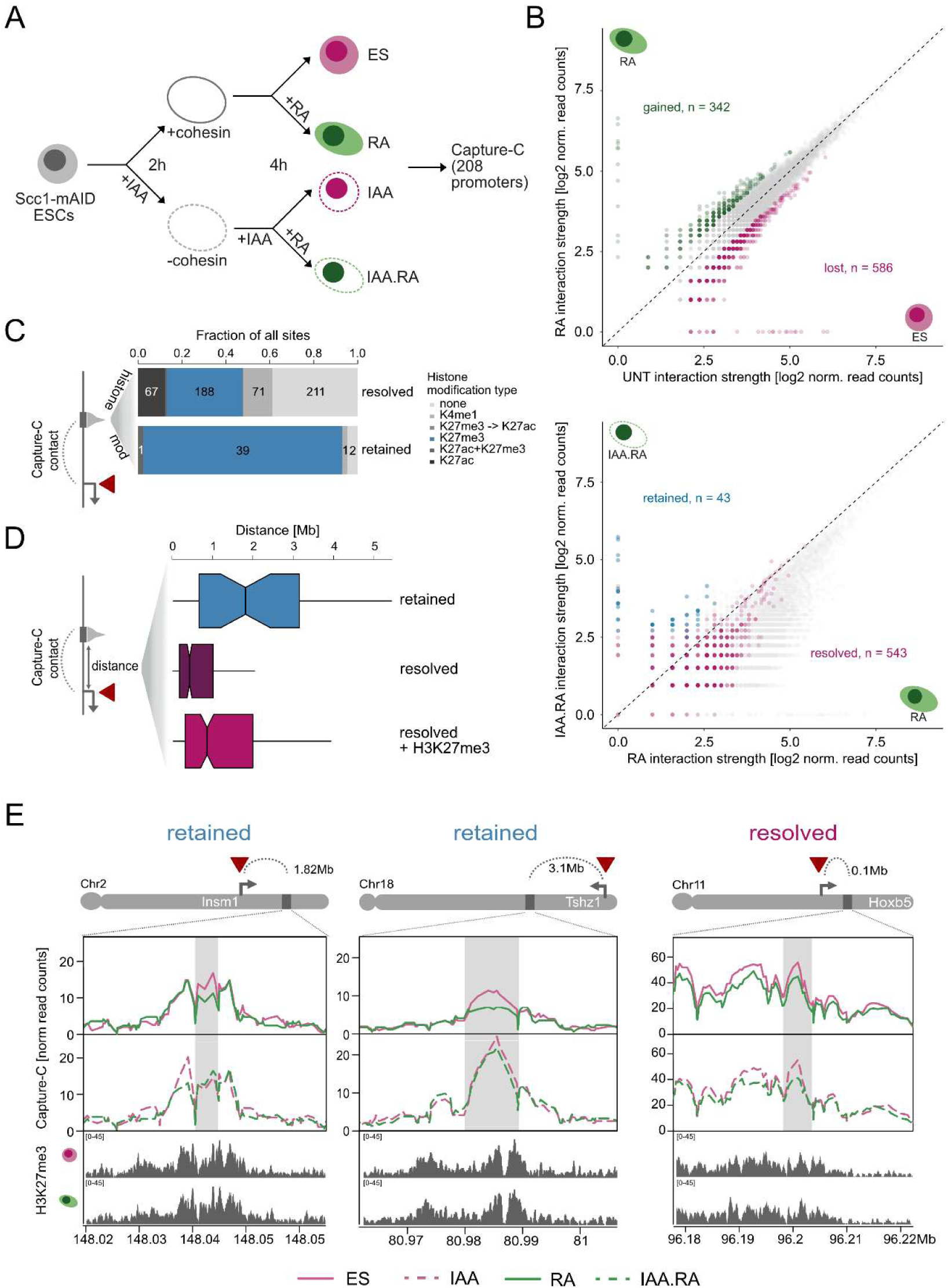
Cohesin is required for the efficient loss of a subset of promoter interactions during embryonic stem cell differentiation. **(A)** Experimental setup. **(B)** Scatterplots showing dynamic promoter interactions in differentiating cells (top) and a comparison between normal and Scc1-depleted differentiated cells (bottom). Interactions lost (red) or gained (green) in normal differentiation are highlighted in the upper plot. Interactions that are lost independently of cohesin (‘resolved’, red) and those that require cohesin for developmental loss (‘retained’, blue) are highlighted in the lower plot. **(C)** The enrichment of histone modifications at distal ends of resolved and retained interactions. **(D)** Comparison of inter-anchor distances between retained and resolved interactions. **(E)** Capture-C lineplots demonstrating promoter interactions upon cohesin loss, including respective H3K72me3 ChIP-seq signal at distal sites^38^. Retained (Insm1 and Tshz1) and resolved (Hoxb5) distal sites are highlighted in grey. Interaction signal has been smoothed over 5 adjacent restriction fragments (k=5). The red triangles in (**C-E**) indicate the location of the capture view point (baits).

### Interactions that depend on cohesin for developmental disruption are enriched for Polycomb-associated histone modifications

To study cohesin’s role in developmental disruption of interactions, we defined a stringent set of reproducibly ‘retained interactions’ (n=43, blue dots in **Fig. 2B**, lower panel) as *cis*-chromosomal interactions that were lost during normal differentiation, but persisted in the absence of cohesin. In contrast, ‘resolved interactions’ (n=543, red dots in **Fig. 2B**, lower panel) are cohesin-independent for their developmental loss. For these two sets of interactions, we next asked how they differed from each other. To determine whether retained interaction anchors vary in their activity compared to sites with resolved interactions, we reanalyzed previously published ChIP-seq data detailing histone modifications before and after RA treatment^38^. This analysis revealed that both promoters and distal sites with retained interactions are highly enriched for H3K27me3-overlapping sites, suggesting that they constitute Polycomb targets (**Fig. 2C, 2E** and **S2D**, empirical p-value < 0.01).

Such association with Polycomb is reminiscent of the Polycomb-associated interactions disrupted by cohesin in ESCs^23^. Similarly to these interactions, that depend on cohesin for disruption in steady-state cells, retained interactions spanned longer distances than resolved interactions, even if those were also associated with Polycomb (**Fig. 2D**).

Some of the retained interactions visually appeared stronger already in ESCs upon cohesin depletion, although this could not be captured in our stringent analysis (**Fig. 2E and S2C**). For the majority, however, cohesin was only acting as a disruptor upon differentiation. Taken together, this suggests that cohesin has a disruptive function during the developmental loss of a subset of long-range interactions that may differ from its disruptive activity in ESCs.

### Retained interactions depend on Polycomb to the same extent as Polycomb-occupied resolved interactions

While H3K27me3 is highly enriched at distal sites with retained interactions, a subset of resolved interactions is also associated with Polycomb (**Fig. 2C**). Seeking to understand why these two sets of Polycomb-occupied sites displayed a different dependency on cohesin for dissolution, we wondered whether retained interactions are more Polycomb-dependent than resolved interactions. To test this, we conducted Capture-C in ESCs in which RING1B, the catalytic subunit of PRC1 complexes, was conditionally depleted^23^ and compared interaction frequencies between resolved and retained interactions (**Fig. 3A**). Strikingly, all Polycomb-associated interactions equally relied on RING1B (**Fig. 3B**), including resolved interactions with promoters that also engaged in retained interactions (**Fig. S2E**). This effect was specific, since those interactions that were not associated with Polycomb remained stable upon RING1B depletion (**Fig. S3A**). Thus, Polycomb can regulate both retained and resolved interactions, suggesting that Polycomb-dependence cannot explain why retained interactions in particular require cohesin for developmental loss.

**Figure 3.**
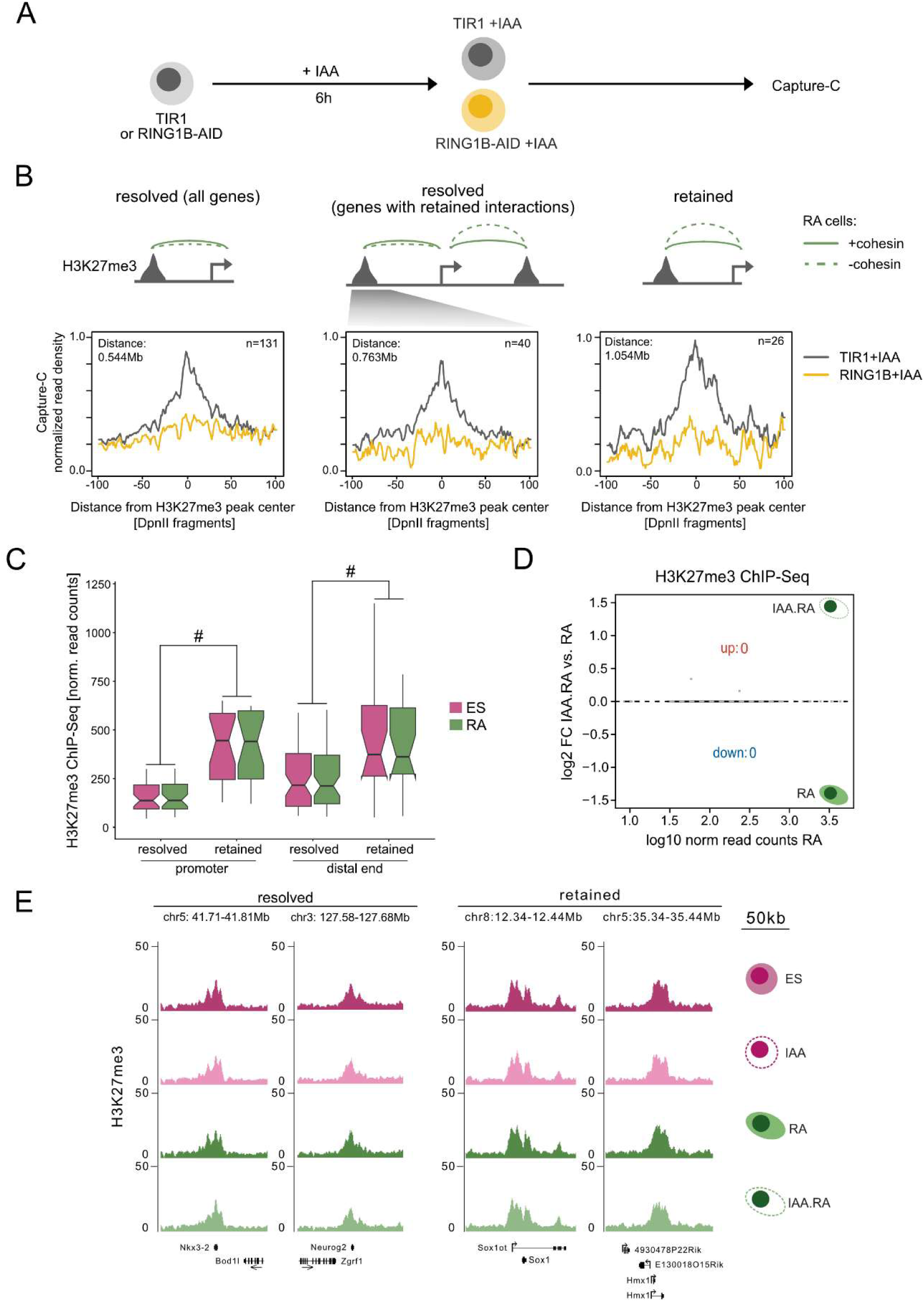
Retained interactions depend on Polycomb to the same extent as Polycomb-occupied resolved interactions. **(A)** Experimental set-up: RING1B-AID cells^23^ were treated with IAA for 6 hours to deplete RING1B. IAA-treated Tir1-expressing parental cell line was used as a negative control. **(B)** Aggregated Capture-C plots demonstrating normalized interaction signal centred at H3K27me3 peaks at resolved and retained sites before (grey) and after (yellow) RING1B depletion. Signal was normalized to the signal at the interaction summit of RING1B-AID. **(C)** Boxplots demonstrating the H3K27me3 enrichment at promoters and distal ends of resolved and retained interactions, as measured by ChIP-seq. Significant comparisons (adjusted p value < 0.05) between resolved and retained loop anchors are indicated by #. All H3K27me3 enrichment comparisons between ES and RA are not significant. **(D)** MA-plot comparison of H3K27me3 levels between cohesin-depleted (IAA.RA) and wild-type (RA) cells (|fold change| ≥ 1.5, padj < 0.05, n=4 replicates). **(E)** H3K27me3 ChIP-seq profiles across all conditions at selected resolved and retained distal sites (smoothing window 5).

Previous studies suggested that interactions with high Polycomb levels are constantly disrupted by cohesin in ESCs^23^, whereas most of the retained interactions appear to only require cohesin for disruption upon differentiation. We therefore speculated that this differentiation-dependent reliance on cohesin may result from a further increase in Polycomb levels at the retained sites during differentiation.

Alternatively, resolved sites may exclusively lose H3K27me3 during the differentiation, thereby losing their ‘glue’. However, a comparison of H3K27me3 levels before and after RA treatment, revealed stable H3K27me3 levels (**Fig. 3C, 3E** and **S3B-C**), suggesting that cohesin dependence of retained interactions for developmental disruption is not a function of elevated Polycomb levels upon differentiation.

Finally, we considered the possibility that H3K27me3 levels are selectively altered by cohesin depletion, rendering retained interactions more cohesin-dependent than resolved interactions. We tested this possibility by comparing H3K27me3 levels before and after IAA treatment, revealing that loss of cohesin was not associated with a loss of H3K27me3 from resolved sites, nor with an increase in H3K27me3 at retained sites (**Fig. 3D-E**). Thus, neither differentiation-dependent nor cohesin-dependent alterations in Polycomb levels can explain the differences between retained and resolved interactions. These results suggest that cohesin is required for programmed developmental loss of long-range Polycomb-dependent interactions in a process that appears independent of dynamic alterations in Polycomb levels.

### Retained interactions are enriched for sequence-specific transcription factor binding

We reasoned that if loss of retained interactions indeed occurs in a programmed manner, it may be associated with the recruitment of specific TFs at these sites. Given the relationship between high cohesin levels and enhancer activity^14^, such recruitment may also explain the cohesin dependence of retained interactions. We therefore set out to test whether a distinguishing feature of retained interaction anchors may be their distinct occupancy by sequence-specific TFs. To increase the statistical power of this analysis, we sought to identify additional retained interactions. Since the same gene can be involved in both retained and resolved interactions (**Fig. S2E**), we conducted reciprocal Capture-C targeting H3K27me3 peaks within 1kb of either resolved (n=78) or retained (n=24) distal sites. This experiment allowed us to validate previously detected retained interactions (**Fig. S4A-C**) and identify nine new retained interactions associated with previously resolved DREs (**Fig. S4C**). To further increase the number of sites in our analysis, we defined ‘retained anchors’ and ‘resolved anchors’ as all H3K27me3 peaks within 1kb of a corresponding site (promoter or DRE) in either the original or the reciprocal Capture-C, thus resulting in 64 retained and 198 resolved Polycomb-associated interaction anchors in total.

Using this expanded set of retained and resolved interaction anchors, we tested their differential TF occupancy by comparative ATAC-footprinting analysis (**Fig. 4A**). Similar to other sites with lost interactions (**Fig. S1B**), accessibility remained stable at both types of anchors during the differentiation (**Fig. S5A**). However, aggregated ATAC-seq peak profiles showed that anchors with retained interactions displayed a generally higher accessibility than anchors with resolved interactions, potentially indicating a more active regulatory state (**Fig. S5A**). Quantitative comparison of TF footprints associated with retained and resolved interaction anchors revealed that 51 and 47 TF motif footprints were enriched at retained sites before and after the differentiation, respectively (**Fig. 4B and S5B**). Retained-anchor enriched TF motifs constituted a diverse group, and included for instance zinc-finger proteins (such as ZNF354A, ZNF143 and ZNF547), differentiation-associated transcription factors (such as NRA5A2 and RARB) and transcriptional repressors (such as ERF). Conversely, only one TF motif (RFX7) was enriched at resolved interaction anchors upon differentiation, indicating that retained interactions are controlled by a distinct set of TF binding motifs. 36 of the retained-enriched TF motifs were enriched at both time points (**Fig. S5C**), with time-point specific enrichment mildly associating with corresponding transcriptional changes (**Fig. S5D**).

**Figure 4.**
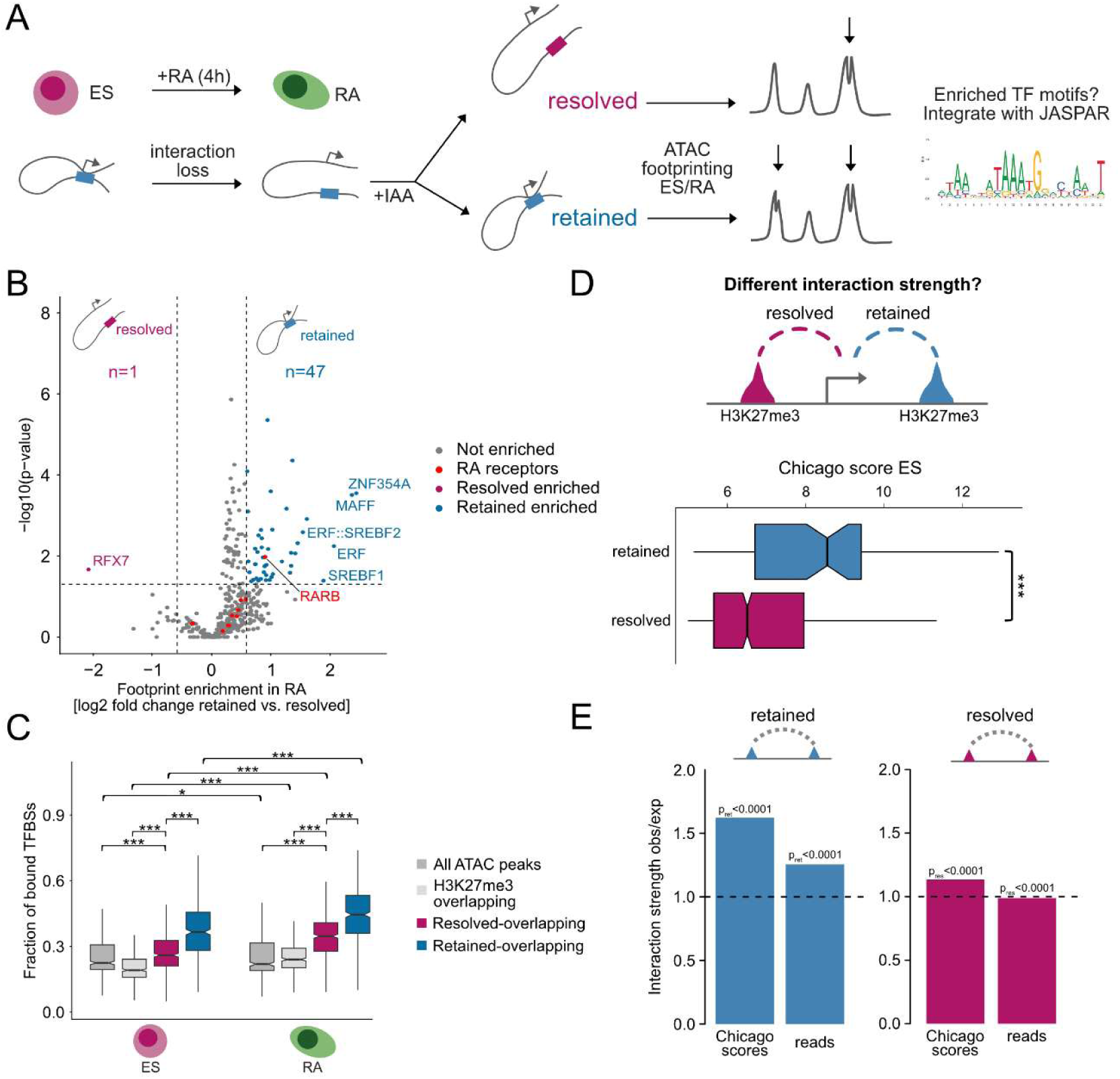
Retained interactions are enriched for sequence-specific transcription factor binding. **(A)** Comparative ATAC-seq footprinting analysis was performed before (ES) and after (RA) the differentiation. **(B)** Volcano plot demonstrating TF footprint enrichment at retained compared to resolved sites after 4h of RA treatment. Top five TFs with the highest enrichment and the enriched RARB are labelled. **(C)** Boxplots show the fraction of bound TFBSs at retained (blue) and resolved (red) interaction anchors compared to ATAC and H3K27me3 peaks at both time points. **(D)** Boxplots comparing the interaction strength of retained (blue) and resolved (red) interactions. **(C-D)** Adjusted p values are shown when significant: ∗p < 0.05, ∗∗∗p < 0.001. **(E)** Barplots depicting observed/expected ratios of interaction strength for retained (blue) and resolved (red) interactions. Expected interaction strengths were derived from a sampling of distance-matched control interactions.

Consistent with higher accessibility, both the fraction and the total number of bound TF motifs, were elevated at retained interaction anchors (**Fig. 4C** and **S5E**). Such high total occupied motif numbers may be explained by the larger size of the retained-anchor associated Polycomb peaks, and further by their closer proximity to TSSs (median values: 2,645 bp and 6,599 bp, for retained and resolved anchors, respectively, as identified in the original Capture-C). The already high TF motif occupancy at retained sites increased even further upon differentiation (**Fig. S5F and 4C**), as did the occupancy of other Polycomb-occupied interaction anchors, but not of ATAC-seq peaks in general (**Fig. 4C**). Among the TF motifs with retained-anchor specific increased occupancy, we detected the binding motif of the retinoic acid receptor RARB, whose expression was also induced during the RA-mediated differentiation (**Fig. 4B, S5B** and **S5F**). This may indicate a specialised response to RA signalling at these sites. Despite this general increase, most TF motifs only displayed moderate changes in their occupancy over time. Indeed, comparative footprinting analysis detected one TF motif with significantly increased footprinting for retained (THRA) and one for resolved (ZFP809) interaction anchors (**Fig. S5G**). Thus, retained interaction anchors are not only enriched for Polycomb, but also display a high accessibility and TF levels.

Given their association with genome organization^24,25,28–34^, we next wondered whether such high occupancy by both TFs and Polycomb is reflected in particularly strong interactions at retained sites. Indeed, retained interactions had higher interaction scores than resolved interactions (**Fig. 4D**), with both score and read counts significantly exceeding those expected at similar distances (**Fig. 4E**). Therefore, retained interaction anchors are enriched for a specific set of highly occupied TF motifs, whose occupancy further increases in differentiation, and is associated with strong long-range interactions. Together, these data indicate that retained interaction anchors may constitute important regulatory platforms that differ from resolved interaction anchors.

### Retained interaction anchors undergo increased rewiring during differentiation

Having detected increased TF occupancy during the differentiation, especially at retained interaction anchors, we next wondered how these TFs may contribute to the programmed developmental loss of interactions at these sites. Binding of TFs to chromatin has been shown to coincide with increased looping, as well as with increased cohesin recruitment in steady cell states^19,28,30,34,44^. Therefore, we hypothesized that the elevated TF occupancy at retained interaction anchors may trigger an increased cohesin-dependent rewiring upon differentiation (**Fig. 5A**), which could either correspond to a stronger interaction gain, or to an increased number of gained interactions compared to those gained by resolved interaction anchors (**Fig. 5A**). To test which of these hypotheses is true, we used the data from our reciprocal Capture-C. After confirming that interactions remained on average stable during differentiation (**Fig. S6A**), we identified all gained interactions (**Fig. 5B**), and quantitatively compared them based on whether their gain occurred with retained or resolved interaction anchors. This analysis revealed that the magnitude of the interaction gain was indistinguishable between these two sets of sites (**Fig. 5C**). Moreover, retained and resolved anchors gained interactions with similar regulatory elements (**Fig. S6B**). Thereby the rewiring of retained and resolved interaction anchors equally depended on cohesin (**Fig. 5C**), and this dependence was not a consequence of altered activity at distal sites, since both H3K27ac and H3K27me3 remained stable in the absence of cohesin (**Fig. S6C and S3B-C**). In sum, cohesin is required for the differentiation-dependent rewiring of all interactions, and retained interaction anchors are not particularly dependent on its function for developmental gain of interactions.

**Figure 5.**
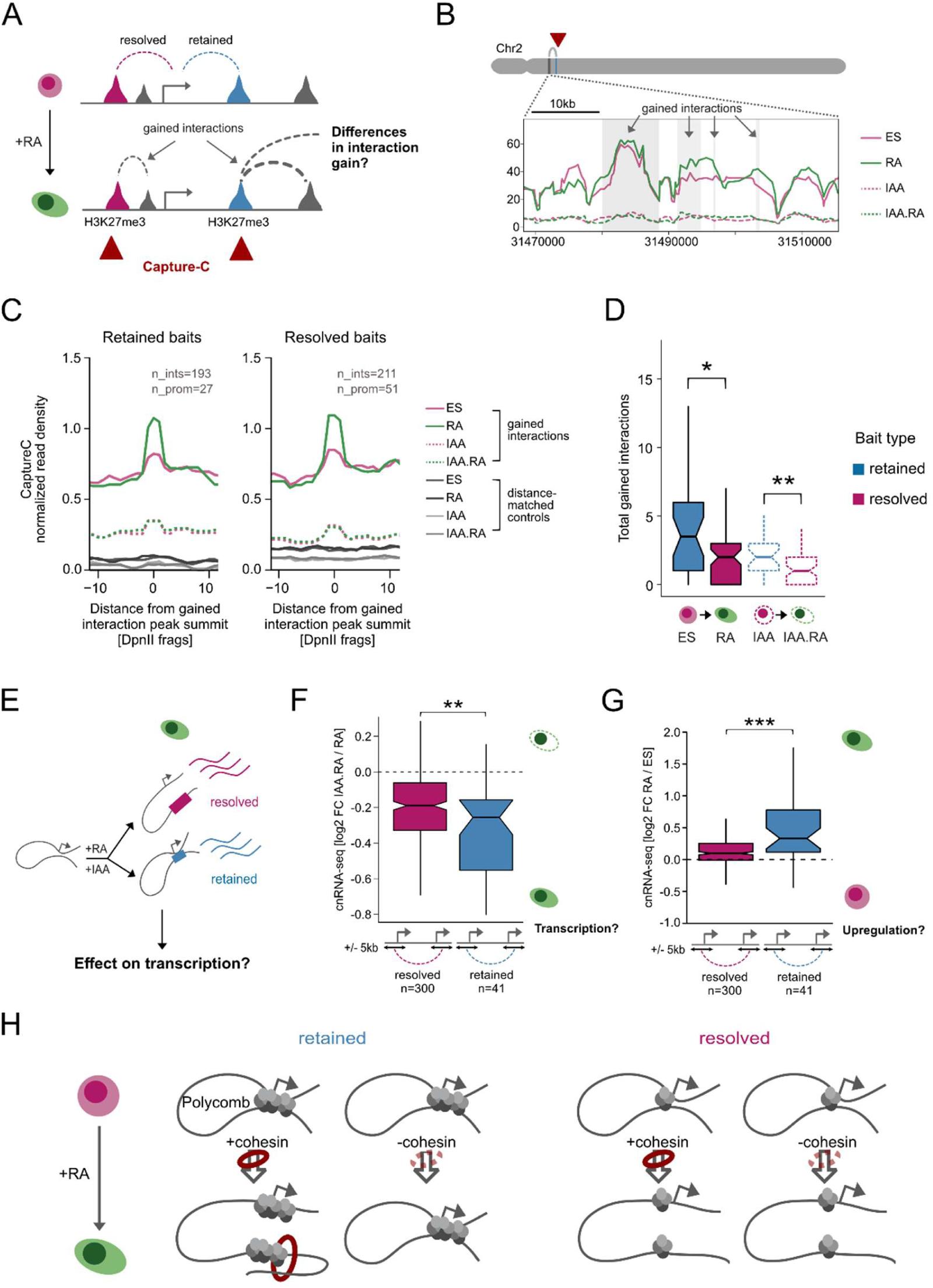
Retained interaction anchors undergo increased rewiring during the differentiation. **(A)** Schematic of the experiment. Distal sites were captured in reciprocal Capture-C to determine their rewiring. **(B)** Example Capture-C snapshot of interactions gained during the four-hour differentiation with a retained interaction anchor. **(C)** Aggregate Capture-C profiles of developmentally gained interactions centred at gained interaction peak summits captured by either retained (left) or resolved (right) baits. Signal was normalized to the signal at the interaction summit in ES and smoothed over k=3 restriction fragments. **(D)** Boxplots displaying the total number of interactions gained by retained and resolved distal sites with and without (dashed lines) cohesin. **(E)** Schematic of the comparison of transcription between genes associated with either retained or resolved interactions. Only genes expressed in RA are considered. **(F-G)** Boxplots showing transcriptional changes upon either cohesin depletion **(F)** or differentiation **(G)** of expressed genes located within 5kb of either resolved (red) or retained (blue) interaction anchors based on cnRNA-seq^38^. **(D-G)** Adjusted p values are shown when significant: ∗p < 0.05, ∗∗p < 0.01, ∗∗∗p < 0.001. **(H)** Schematics demonstrating how Polycomb-mediated interactions rewire during differentiation, and how this process is affected upon cohesin loss.

In contrast to the magnitude of interaction gain, retained interaction anchors acquired significantly more interactions during the differentiation than resolved interaction anchors, indicating a more active rewiring (**Fig. 5D**). We noted that sites with retained interactions remained promiscuous even in the absence of cohesin, acquiring a total of 98 interactions, only 9 of which overlapped with those gained upon the differentiation of wild-type cells (**Fig. 5D and S6A**). Similar to retained interactions, these ectopically acquired interactions were enriched for Polycomb-associated histone modifications (**Fig. S6B**) and reduced during normal differentiation, as visible in the aggregate peak analysis (**Fig. S6A**). However, in contrast to retained interactions, they still remained weaker in cohesin-depleted cells compared to wildtype cells and thus were not identified as retained interactions. This suggests that in normal differentiation cohesin counteracts the acquisition of putatively repressive interactions, and instead ensures that retained interaction anchors undergo a strong rewiring characterized by both cohesin-dependent gain and loss of interactions.

We reasoned that this strong rewiring may be related to the general properties of differentiation-dependent TAD restructuring. This implies that the developmental separation of interaction anchors occurs due to their partitioning from the same TAD into two separate TADs upon RA treatment. We therefore compared how TAD identity changes upon differentiation for resolved and retained interactions, using published Hi-C data from ESCs and neuronal progenitors^40^. Consistent with their supra-megabase long distances from each other, anchors of retained interactions were more likely to reside within different TADs in ESCs than the anchors of resolved Polycomb-associated interactions (90% vs 70%). Strikingly, their partitioning into separate TADs did not increase upon differentiation (**Fig. S6D**). Therefore, the cohesin-dependency of retained interactions is unlikely to occur due to increased TAD partitioning upon differentiation.

Together, these data indicate that Polycomb-occupied sites with retained interactions differ from those with resolved interactions in their enhanced active rewiring upon RA-mediated differentiation.

### Genes associated with retained interactions are transcriptionally downregulated

In order to understand the significance of this cohesin-dependent rewiring of interactions, we next wondered how it may affect gene expression. We reasoned that in the absence of cohesin, retained interaction anchors maintain their interactions with potentially repressive Polycomb sites, while simultaneously failing to acquire potentially activating interactions. In contrast, genes with resolved interactions do not gain activating interactions without cohesin, but also do not maintain interactions with repressive sites. If loss and gain of interactions are both important for transcription, then the dual effect of excessive repressive and insufficient activating signal may lead to a stronger transcriptional downregulation of the genes associated with retained interaction anchors. To test this, we used our previously published data^38^ in which we first selected all RA-expressed transcripts whose TSSs were located within 5 kb of a retained or of a resolved interaction anchor (**Fig. 5E-G**). The distance of 5 kb was chosen to accommodate the average size of H3K27me3 peaks at these anchors (9.3 kb). For the resulting 41 transcripts near retained and 300 near resolved interaction anchors, we compared the effect of cohesin depletion on their expression upon differentiation. This revealed that retained-anchored genes were more downregulated in the absence of cohesin than genes near resolved interaction anchors (**Fig. 5F**). Of note, genes with resolved interactions were also transcriptionally downregulated, possibly due to impaired acquisition of potentially activating interactions. In addition to being disproportionately affected by cohesin depletion, retained-anchored genes displayed lower expression in ESCs (**Fig. S6E**) and underwent a stronger transcriptional upregulation upon RA-treatment (**Fig. 5G**), suggesting that they constitute developmentally upregulated genes. These data indicate that appropriate differentiation-dependent rewiring, including appropriate programmed loss of interactions, may be important for the transcriptional regulation during embryonic stem cell differentiation.

## Discussion

Cell type transitions are characterized by a widespread loss of long-range interactions, yet it remains unclear how this loss is regulated at the molecular level. Using footprinting analysis, we show that promoter interactions resolve independently of accessibility changes or loss of sequence-specific TF motif occupancy (**Fig. 1**). Instead, we observe that a full developmentally programmed loss of interactions requires cohesin (**Fig. 2**). This requirement is not ubiquitous, but limited to strong long-distance interactions with high Polycomb occupancy. Similar to a subset of cohesin-independently resolved interactions, interactions that are retained in the absence of cohesin are regulated by Polycomb (**Fig. 3**). However, their key difference from Polycomb-occupied resolved interactions lies in increased numbers of sequence-specific TF binding (**Fig. 4**) and a strong, differentiation-and cohesin-dependent rewiring that appears important for appropriate gene expression upon differentiation (**Fig. 5**).

Our findings align with a model in which Polycomb-associated interactions are resolved during ESC differentiation in a programmed manner that involves cohesin, to enable appropriate transcriptional activation of the associated genes. Such programmed developmental disruption may be necessary because of the repressive function of these interactions^45^. However, an equally possible-and not mutually exclusive - scenario is that appropriate regulation of these genes relies on their ability to acquire new activating interactions, as has been proposed by previous studies focusing on cohesin’s role in contact gain^41,46,47^. In support of this possibility, genes that resolved their interactions independently of cohesin, but still failed to acquire new interactions, also are transcriptionally affected by its loss (**Fig. 5**). This effect, however, was smaller than the effect observed for retained genes, suggesting that both gain of activating and loss of repressive interactions may cooperatively regulate transcription during cell type transitions. Interestingly, cohesin appears to modulate both of these processes, acting as a loop-promoting factor, according to its established role^6–9^, while simultaneously contributing to the programmed loss of interactions. This dual activity could provide a mechanistic explanation for the recently reported increased transcriptional dependence of Polycomb-occupied genes on cohesin during the differentiation of dendritic cells^48^.

We propose that efficient dissolution of strong Polycomb-mediated interactions is mechanistically regulated by cohesin-dependent interaction gain, resulting in a simple model, in which cohesin actively rewires interaction anchors by guiding them towards other regulatory sites (**Fig. 5H)**. A side effect of active rewiring is passive disruption of very strong interactions, which we show occurs independently of TAD partitioning, yet may be facilitated by the differentiation-dependent strengthening of TADs^40^. This model aligns with the key role of cohesin as a facilitator of interactions^6–9^ and further extends its proposed role as a disruptor of very long-range interactions in steady cell states^7,22,23^ towards dynamic processes. Nevertheless, we cannot exclude other possible scenarios, which may coexist with this cohesin-dependent rewiring model. For instance, cohesin may act as an active disruptor of promoter interactions, or their loss may be regulated by other DNA binding factors, including those TFs that increase their occupancy upon differentiation^30,31^.

At the moment, we can only speculate how increased rewiring is regulated at retained interaction anchors. It is possible to envision that a causal relationship between TF occupancy and promoter interactions contributes to developmentally programmed interaction loss. For instance, high levels of TF binding at retained sites (**Fig. 4**) may be directly related to the increased strength of their interactions, whilst increased occupancy upon differentiation may actively trigger their rewiring. Since only a small subset of sequence-specific TFs have been directly implicated in long-range interactions, this activity may be mediated either by individual TFs enriched at these sites, or by an ensemble of TFs, reminiscent of the models proposed for enhancer regulation^52^. Thereby, interactions can be regulated both directly or indirectly via the recruitment of additional effectors^14,24–27,53,54^, or condensate formation^54–57^.

A surprising finding of our study is that only a subset of Polycomb-dependent (‘retained’) interactions require cohesin for programmed developmental disruption (**Fig. 2-3**). While this dependency cannot be explained by a differential regulation of H3K27me3, we note that it remains possible that PRC1 levels are selectively altered in the absence of strong H3K27me3 changes^40^. However, given previous observations, we would expect quantitative effects on the H3K27me3 levels within a short period of time^51^. Given that Polycomb occupancy is causally associated with retained interactions (**Fig. 3**), their particular strength in ESCs may be determined by high Polycomb levels, high TF occupancy, or a combination of both. A shift in this dependency during differentiation would allow the differentiation-dependent rewiring to be driven by sequence-specific TFs whose occupancy increases upon differentiation. Mechanistically, these TFs may directly regulate the rewiring of interactions, including both gain and loss of contacts. Alternatively, they could stall cohesin, as has been shown for active RNAP2^15^, thereby enlisting the complex in rewiring of the associated sites. Importantly, cohesin recruitment could occur directly or indirectly, for instance via the recruitment of Mediator complexes^58^.

Finally, our data indicate that a simple loss of sequence-specific factors is insufficient to explain developmental loss of interactions, yet the majority of interaction loss occurs independently of cohesin. This suggests that other factors not included in our analyses are involved in the active or passive disruption of cohesin-independently resolved interactions. These may include the footprint-gaining TF TP53 (**Fig. 1B**), or the resolved-interaction enriched TF RFX7 (**Fig. 4B**). Other factors without a sequence-specific binding domain could also play a role, such as JMJD2^54^, Mediator^26,27^, histone modifying complexes^53^, or RB that has been recently reported to remove cohesin from chromatin^59^. Future studies addressing the role of sequence-specific and unspecific factors in promoting and disrupting interactions via targeted mapping^54,60^, perturbation assays or targeted recruitment will be required to clarify underlying molecular mechanisms.

### Limitations of the study

A limitation of the study is that we are unable to distinguish whether cohesin actively disrupts interactions, or passively regulates interaction loss through redirection of loop anchors towards other sites. Moreover, given that cohesin degradation affects both loss and gain of interactions, it remains to be shown whether resolving interactions is required for full gene expression. More targeted experiments will be required to address both of these points, such as targeted manipulation of looping^49,50^. Another limitation is that we focused on the early differentiation time points (up to 4 hours) to understand the regulatory mechanisms underlying the first wave of interaction loss, which encompasses the majority of developmentally disrupted interactions. With this approach we could not assess the mechanisms underlying the second wave of interaction loss that occurs between 6 and 24h, nor the effect of early interaction loss on genes that are induced later in differentiation^38^. Additional experiments, such as cohesin wash-out^7^, or alternative strategies for manipulation of its loop extrusion activity^47^, will be required to extend the analysis to later time points.

## Acknowledgements

We would like to thank Ralph Grand, Magdalena Laugsch and the members of the Feldmann lab for providing critical comments on the manuscript, Karsten Rippe for productive discussions of the project. We are grateful to the DKFZ Genomics (NGS) and ODCF core facilities for sequencing and data pre-processing support. We are grateful to Martina Capriati and Ralph Grand for providing advice on ATAC-seq, to Jim Hughes, James Davies and Damien Downes for providing advice and protocols for Capture-C prior to publication. Ann-Kristin Dicke is supported by a postdoctoral fellowship from the Dr. Rurainski Foundation. Work in the Feldmann lab is supported by the Helmholtz Young Investigator Award (VH-NG-1604), ERC Starting Grant (ERC-STG 101115812) and Health + Life Science Alliance Heidelberg-Mannheim (Explore!Tech projects ‘delMosaic-seq’ and MWK32-7531-26/7/2, Z02).

## Author Contributions

Conceptualization, V.S. and A.F.; Methodology, V.S., A.K.D. and A.F.; Investigation, V.S., A.K.D. and A.F.; Formal Analysis, V.S., A.K.D. and A.F.; Resources, V.S., A.K.D. and A.F.; Writing – Original Draft, V.S. and A.F.; Writing – Review & Editing, V.S., A.K.D. and A.F.; Funding Acquisition, A.F.; Supervision, A.F.

## Materials and Methods

### RESOURCE AVAILABILITY

#### Lead contact

Further information and requests for resources and reagents should be directed to and will be fulfilled by the lead contact, Angelika Feldmann.

#### Materials availability

This study did not generate new unique reagents.

#### Data and Code Availability

• The high-throughput data reported in this study have been deposited on GEO and are publicly available as of the date of publication. Re-analyzed datasets include Capture-C (GSE261064), ChIP-seq (GSE261039) and RNA-seq (GSE261066) from Mahara et al.^38^

• All original code has been deposited on Zenodo.

• Any additional information required to re-analyze the data reported in this paper is available from the lead contact upon request.

#### EXPERIMENTAL MODEL AND STUDY PARTICIPANT DETAILS

Male E14 mouse ESCs, as well as E14-derived Scc1-mAID cell line^23^ were cultured in high glucose DMEM (Gibco) supplemented with 15% fetal bovine serum (Sigma-Aldrich), 2mM Glutamax, 1x non-essential aminoacids, 1x peniciliin-streptomycin, 0.05mM beta-mercaptoethanol (Gibco) and Leukaemia Inhibitory Factor (LIF, produced in-house). Cells were plated on 0.1%-gelatine coated plates and maintained at 37°C within an incubator with 5% CO_2_. Cells were passaged every two days, and at least two passages prior to processing.

To perform cohesin degradation, 6 x 10^6^ E14-derived Scc1-mAID cells^23^ were plated on 0.1% gelatine-coated 15cm plates. After approximately 12 hours the medium was replaced with medium containing either 500µM indole-3-acetic acid (Merck, #I3750-5G-A) or ethanol. Two hours after treatment initiation, the medium was replaced with EC-10 medium (DMEM, 10% fetal bovine serum, 2mM Glutamax, 1x non-essential aminoacids, 1x peniciliin-streptomycin, 0.05mM beta-mercaptoethanol) containing 1µM retinoic acid (Merck, R2625) and 500µM indole-3-acetic acid for the next four hours. Afterwards, the cells were collected for downstream analyses using TrypLE (Gibco).

HEK293T used for nChIP spike-in were cultured in DMEM supplemented with 10% fetal bovine serum, 2mM Glutamax, 1x non-essential amino acids, 1x penicillin-streptomycin and 0.05mM beta-mercaptoethanol and passaged once the confluency was reached.

### METHOD DETAILS

#### ATAC-seq library preparation

To assess chromatin accessibility before and after 4 hours of RA differentiation, ATAC seq of four biological replicates was performed. Library preparation was carried out using the “Pre-indexed Assembled Tn5 Transposomes” and “ATAC-seq Buffer Set” kits by Active Motif according to the manufacturer’s instructions. Briefly, 100,000 E14 wildtype and 100,000 RA-differentiated cells were obtained and lysed using the ATAC lysis buffer for 10min, followed by centrifugation. DNA was incubated with hyperactive pre-indexed Tn5 transposase for 40 minutes. Sequencing adapters were subsequently added to accessible regions of chromatin via PCR. A purification was carried out using Ampure XP beads (ratio: 1.2x). After pooling, a double-sided size selection, again using Ampure XP beads (ratios 0.6x and 0.9x), followed. Libraries were quantified using Qubit and TapeStation. Sequencing was performed on an Ilumina NovaSeq X+ system, resulting in paired-end 50 bp sequencing reads.

#### Capture-C

Chromatin was extracted and fixed as described previously^38^. Briefly, 10 x 10^6^ mouse ESCs were trypsinized, collected in 50ml falcon tubes, resuspended in 9.3ml medium and crosslinked with 1.25ml 16% formaldehyde (1.89% final; Thermo Fisher Scientific #10751395) while rotating for 10min at room temperature. The fixation was quenched with 1.5ml 1M cold glycine, washed with cold PBS, and lysed for 20min at 4°C in lysis buffer (10mM Tris pH 8, 10mM NaCl, 0.2% NP-40, supplemented with cOmplete protease inhibitors (Roche)). Next, samples were snap frozen in 1ml lysis buffer on dry ice. Fixed chromatin was stored at-80°C.

Capture-C libraries were prepared as described previously^38^. Briefly, lysates of 10 x 10^6^ cells were thawed on ice, pelleted, and resuspended in 650µl 1x DpnII buffer (NEB #B0543). Three 1.5ml tubes with 200µl lysate each were treated in parallel with SDS (Thermo Fisher Scientific #AM9820, 0.28% final concentration, 1h, 37°C, interval shaking 500rpm, 30 sec on/off), quenched with Triton X-100 (1.67% final concentration, 1h, 37°C, interval shaking 500rpm, 30 sec on/off) and digested for 24 hours with 3 x 10µl DpnII (homemade, 37°C, interval shaking 500rpm, 30 sec on/off). 100µl of each aliquot were taken for digestion control, reverse crosslinked and visualized on an agarose gel. The remaining chromatin was then independently ligated with 8µl T4 Ligase (240U, Thermo Fisher Scientific, #EL0013) in a volume of 1440µl (20h, 16°C). Following this, the nuclei containing ligated chromatin were pelleted to remove any non-nuclear chromatin, reverse-crosslinked, and the ligated DNA was phenol-chloroform purified. The sample was resuspended in 300µl water and sonicated 13x (Bioruptor Pico, 30sec on/off) or until a fragment size of approximately 200bp was reached. Fragments were size selected using Ampure XP beads (Beckman Coulter #A63881, selection ratios: 0.85x / 0.4x). Two reactions of 1-5µg DNA each were adaptor-ligated and indexed using the NEBNext End Repair, dA-Tailing and Quick Ligation modules (NEB #E6050, #E6053, #E6056) and NEBNext Multiplex Oligos for Illumina Primer sets 1 (NEB #E7335) and 2 (NEB #E7500S). The libraries were amplified with 7 PCR cycles using Herculase II Fusion Polymerase kit (Agilent #600677). Libraries were next hybridized in the following way: For each promoter-containing DpnII restriction fragment we designed two capture probes (aligned to mm10, 70 or 120bp each) using the online tool by the Hughes lab (CapSequm: https://oligo.readthedocs.io/en/latest/)^61^ with the following filtering parameters: Duplicates: <2, Density <30, SRepeatLength <30, Duplication: FALSE. For promoters for which no probes could be designed for the restriction fragment directly overlapping the TSS, probes were designed for the next-nearest DpnII fragment, if it was within 500bp of the TSS. The probes were pooled at 2.9nM (RING1B Capture-C) or 0.2nM (reciprocal Capture-C) each and the samples were multiplexed by mass prior to hybridization (1-2µg each, according to Qubit dsDNA BR Assay, Thermo Fisher Scientific #Q32850). Hybridization was carried out using the KAPA HyperCapture Reagent Kit with KAPA HyperCapture Bead Kit (Roche #9075828001 and #9075798001) according to Roche protocol for 72 hours, followed by a second 24-hour hybridization step (double Capture). Captured libraries were quantified by qPCR using the KAPA Sybr Fast Universal kit (KAPA #KK4602) and sequenced on Illumina NextSeq 550 or Illumina NovaSeq X+.

#### Calibrated native ChIP library preparation

Calibrated native ChIP-seq to map H3K27me3 and H3K27ac was performed as described previously with minor changes^62^. Briefly, 25 × 10^6^ ES cells were mixed with 1 × 10^6^ HEK293T cells, washed with 1× PBS and resuspended in 1ml ice-cold RSB buffer (10mM Tris-HCl pH 8.0, 10mM NaCl, 3mM MgCl2) prior to chromatin isolation. Nuclei were released by adding 14ml of ice-cold lysis buffer (10mM Tris-HCl pH 8.0, 10mM NaCl, 3mM MgCl2, 0.1% NP-40, 5mM sodium butyrate). Nuclei were subsequently washed, isolated by centrifugation at 1500g for 5min at 4°C, resuspended in ice-cold sucrose-calcium chloride solution (10mM Tris-HCl pH 8.0, 10mM NaCl, 3mM MgCl2, 0.25M sucrose, 3mM CaCl2, 5mM sodium butyrate, 1× cOmplete protease inhibitor cocktail (Roche)) and treated with 100 units of micrococcal nuclease (Thermo Fisher Scientific #EN0181) for 5min at 37 °C. The digestion was stopped with the addition of 4µl 0.5M EDTA. The samples were centrifuged at 1500g for 5min at 4°C and the supernatant containing digested chromatin fragments (S1) was harvested. The remaining pellet contained chromatin fragments trapped in the nucleus, which were released by incubation with 300μl of nucleosome release buffer (10mM Tris-HCl pH 7.5, 10mM NaCl, 0.2mM EDTA, 5mM sodium butyrate, 1× protease inhibitor cocktail (Roche)) at 4°C for 1h, passed through a 27-gauge needle five times, and centrifuged at 1500g for 5min at 4°C. The second supernatant (S2) was harvested and combined with S1 supernatant, and the purified native chromatin fragments were aliquoted into dry ice pre-chilled tubes (∼5 × 10^6^ ES cells/tube) and stored at-80 °C. Nuclease digestion was assessed with a 1.5% agarose gel.

For a single ChIP, purified nuclease-digested chromatin from 5 × 10^6^ ES cells aliquot was diluted 10-fold with the native ChIP incubation buffer (10mM Tris-HCl pH 7.5, 70mM NaCl, 2mM MgCl2, 0.1% Triton X-100, 2mM EDTA, 5mM Sodium butyrate, 1× cOmplete protease inhibitor cocktail (Roche)). Next, the chromatin was pre-cleared for 2h with Protein A Dynabeads (Thermo Fisher Scientific #10001D) blocked with 0.05mg/ml tRNA and 0.2mg/ml BSA. A 10% aliquot of precleared chromatin was taken as input, while the rest was further incubated overnight at 4°C with the appropriate antibody: anti-H3K27ac (CST, #8173, 5µg), anti-H3K27me3 (Millipore, #07-449, 6µg). Antibody-bound chromatin was captured with blocked Dynabeads for 1h at 4°C and isolated through magnetizing on ice. After four washes with ice-cold native ChIP wash buffer (20mM Tris-HCl pH 7.5, 125mM NaCl, 0.1% Triton X-100, 2mM EDTA) and one wash with ice-cold 1x TE buffer, the ChIP DNA was eluted in fresh elution buffer (1% SDS, 0.1 M NaHCO3) via shaking at 25°C for 30min. Both ChIP and input DNA were purified using ChIP DNA Clean and Concentrator Kit (Zymo Research #D5205) following the manufacturer’s protocol.

The sequencing libraries were prepared in accordance with the protocol for NEBnext Ultra II DNA library preparation kit (#E7645). Libraries were quantified using Qubit and Bioanalyzer and sequenced with Illumina NextSeq550 to obtain 42bp paired reads.

#### Whole cell nuclear extraction and Western blot

Collected cells were lysed with RIPA lysis buffer (20mM Tris-HCl pH 7.5, 150mM NaCl, 1% NP-40, 0.5% sodium deoxycholate, 1mM EDTA, 0.1% SDS, 1× cOmplete protease inhibitor cocktail (Roche)) for 1 hour on ice, vortexing every 15min. Next, the lysates were centrifuged at 13,000g for 5min at 4°C, and the supernatant containing whole cell lysate was collected. The protein concentration was measured with the Bradford assay using Protein Assay Dye Reagent Concentrate 5x (BioRad #5000006).

Then, 20μg of protein extract was mixed with 1× SDS loading buffer (2% SDS, 0.1M Tris pH 6.8, 0.1M DTT, 10% glycerol, 0.1% bromophenol blue) and placed at 95°C for 5min. The sample was electrophoretically resolved using home-made 8% SDS-PAGE gels and further transferred onto nitrocellulose membranes using the Trans-Blot Turbo transfer system (BioRad #17001918) according to the manufacturer protocol. Following blocking and incubation with the appropriate primary and secondary antibodies, the membranes were washed two times with 1x PBS, 0.1% Tween-20 and once with 1x PBS, and developed using the Clarity Western ECL Substrate (BioRad #1705060). The chemiluminescent signals were detected using Chemidoc imaging system (BioRad). Antibodies used for Western blot analysis were diluted in 5% w/v non-fat dry milk, 1x PBS, 0.1% Tween-20: rabbit polyclonal anti-RAD21 (Abcam, #AB154769, 1:2000), rabbit monoclonal anti-TBP (Abcam, #ab220788, 1:2000), donkey HRP-linked anti-Rabbit IgG (Cytiva, #NA934, 1:5000).

### QUANTIFICATION AND STATISTICAL ANALYSIS

#### ATAC-seq data processing

Paired-end FASTQ files were merged per sample and quality was assessed using FastQC v0.12.1 (http://www.bioinformatics.babraham.ac.uk/projects/fastqc). Reads were aligned to the mouse reference genome (mm10) using Bowtie2 v2.5.4^63^. Alignments were sorted and PCR duplicates were removed using SAMtools v1.20^64^ and Sambamba v1.0.1^65^. Reads overlapping ENCODE blacklisted regions^66^ were filtered out. Mapping statistics were assessed using SAMtools flagstat, and only properly paired reads were retained for downstream analyses. Quality control across replicates was performed using deepTools v3.5.1 (high correlation between replicates, spearman correlation > 0.96)^67^. Peak calling was carried out using the MACS2^68^ callpeak function with the --broad option and genome size parameter-g mm. Peak calling parameters were set to-q 0.01 and --broad-cutoff 0.05. Peaks were called individually for each replicate. Consensus peak sets per condition were generated by merging replicate peak intervals using the reduce() function from GenomicRanges^69^, retaining only those peaks present in at least 3 out of 4 biological replicates. To control for differences in sequencing depth, filtered BAM files were downsampled to the smallest library size using SAMtools-based random subsampling. Downsampled replicates were re-merged per condition, coverage tracks were regenerated, and peak calling was repeated using the same MACS2 parameters. Consensus peaks for downsampled data were defined analogously (≥3/4 replicates) and merged to obtain final high-confidence peak sets.

#### Differential accessibility analysis using DESeq2

Differential accessibility analysis was performed using DESeq2^70^ on read counts quantified over the merged consensus peak set. For differential analysis, low-abundance peaks (row sum ≤3) were filtered out. A DESeqDataSet was constructed with a design formula including replicate and condition effects to account for batch effects between biological replicates. The baseline condition was set to 0h (wildtype E14 cells). Wald tests were performed for 4h compared to 0h, and log2 fold changes were shrunk using lfcShrink. Peaks were considered significantly differentially accessible at adjusted p-value (padj) < 0.05 and |LFC| ≥ log2(1.5). MA plots were generated, highlighting significantly up-and downregulated peaks. In addition, genomic regions that lose interactions within the first four hours of RA differentiation were intersected with peak coordinates using GenomicRanges, and overlapping peaks were highlighted in MA plots.

#### ATAC footprinting analysis

Transcription factor footprinting analysis was performed using the TOBIAS tool^71^. BAM files were first bias-corrected using ATACorrect, and footprint scores were computed genome-wide using ScoreBigwig. Differential transcription factor binding of 0h and 4h RA differentiation samples was assessed using BINDetect in time-series mode with mm10 as reference genome and merged consensus broad ATAC peaks as the peak set. Motif screening was conducted using curated MEME-format motif libraries obtained from the JASPAR database (JASPAR2024 CORE vertebrates collection [non-redundant pfms meme])^72^. BINDetect was run on all ATAC peaks and separately on distinct genomic subsets by specifying different --output-peaks BED files: (i) all H3K27ac-marked regions (control set), (ii) regions losing chromatin interactions within the first 4 hours of RA differentiation, (iii) peaks overlapping with genomic regions with resolved interactions and (iv) with retained interactions in the absence of cohesin. This strategy enabled comparison of transcription factor binding dynamics between these classes. Differential binding was computed by BINDetect across both time points.

#### ATAC Visualization

TOBIAS BINDetect output tables were imported into R 4.5.1 and filtered to keep motifs with ≥5 predicted binding sites and mean FPKM ≥1 at 0h or 4h in RNA seq data. Heterodimer motifs were kept only if both components were expressed. For all ATAC peaks, retained and resolved interaction regions, fraction-based enrichment was calculated between 4h and 0h (bound/total TFBS in subset), as well as enrichment of factors in retained vs. resolved elements, and significance was assessed using Fisher’s exact test. Volcano plots were generated displaying log2 fold enrichment on the x-and –log10(p-value) on the y-axis, highlighting significantly enriched TFs (p < 0.05, fold change ≥1.5) and RA-related TFs in red.

TFs significantly enriched in retained regions were separately extracted for 0h and 4h and compared using Venn diagrams to identify shared and time point–specific factors. RNA expression (FPKM) of TFs belonging to each Venn category (0h only, shared, 4h only) was visualized using boxplots, and differences between 0h and 4h per category were assessed using unpaired Wilcoxon tests (multiple testing correction = Benjamini and Hochberg).

Global TF binding distributions were compared between control region sets (H3K27me3-marked regions and all consensus ATAC peaks), retained and resolved using boxplots. For each TF (≥5 predicted binding sites and expressed at 0h or 4h), binding was quantified at 0h and 4h either as (i) the fraction of bound TFBS relative to total predicted TFBS or (ii) the number of bound TFBSs normalized to the number of intervals within each region class. Paired Wilcoxon signed-rank tests were used for within-class temporal comparisons (0 h vs 4 h), whereas unpaired Wilcoxon rank-sum tests were used for between-class comparisons at each time point, with Benjamini–Hochberg correction applied in both cases. Boxplots with notches indicating median confidence intervals were generated using ggplot2.

For TFs significantly enriched in retained elements, the fraction of bound TFBS was calculated at 0h and 4h within retained and resolved elements. Trajectories were visualized as line plots connecting 0h and 4h measurements for each TF. RARB was specifically highlighted in red.

#### Capture-C analysis

Paired-end reads were aligned to mm10 and filtered for Hi-C artefacts using HiCUP v 0.8.3 ^73^ and Bowtie2 v2.3.5.1^63^, with fragment filter set to 100-800bp. Read counts for reads that align to the captured gene promoters, as well as interaction scores, were then called by the Bioconductor package Chicago v1.6.0 using the following parameters: minFragLen=75, maxFragLen=40000^74^.

Upon inspection of Chicago plots and evaluation of high quality of interactions, all downstream analysis was conducted using the R package PostChicago^75^. Significant interactions were extracted using makeIntsTable() with the following score cutoffs: scorecut=5 for the original and reciprocal Capture-C experiments; scorecut=3 for the RING1B-AID samples (Fig. 3).

Following this, replicates were correlated and inspected for accurate clustering and high correlation of downsampled read counts (>0.8) and scores (>0.55).

#### Capture-C visualization and comparative analysis

Capture-C visualization was conducted similarly to previous publications^23,26,38^. To visualize the Capture-C data in line plots, we used the PostChicago function plotInteractions(). Briefly, the weighted pooled read counts from the Chicago data files were normalized to the total read count aligning to the captured gene promoters in the sample, and then to the number of promoters in the respective capture experiment. This was then multiplied by a constant number to simplify the visualisation, using the following formula: normCounts=1/cov*nprom*100000. Running means across adjacent fragments were plotted with k fragments indicated for each of the plots.

For scatterplots, reads were downsampled based on the read counts within the captured library using the read counts aligned to baited fragments and determined by Chicago using the function makeIntsTable(). Log2 transformation of scores and reads was performed after the addition of a pseudocount of 1. We used the function annotateInts() to determine the types of histone modifications at distal sites, where significant interactions (or aggregated interaction peaks) were overlapped with peak intervals either from Mahara et al.^38^, or from this manuscript with a peak expected to be within 1000bp (or overlapping for aggregated peaks).

For Aggregate Peak analysis in Fig. 5, significant interactions (Chicago score >=5) within an empirically chosen distance of two restriction fragments were aggregated to single interaction peaks using the function reduce() from GenomicRanges. makeOneGeneOnePeak() and getMatrix() were used to create normalized and distance-matched matrices surrounding the centres of either ChIP-seq (Fig. 3) or interaction (Fig. 5) peaks, which were subsequently plotted using plotAggregatePeaks(). Interaction peaks were also used for the quantification of gained interactions upon rewiring in Fig. 5.

#### Definition of lost, gained, retained and resolved interactions

Lost interactions were defined as previously described^38^. Briefly, they were required to be present in ESCs (Chicago score >=5, minimum read count>=3) and absent upon RA-treatment (score <3 for the original and <5 for the reciprocal Capture-C). Total read counts were required to be 1.5x higher in ESCs than upon RA treatment and further required to be higher in ESCs in each individual replicate. Gained interactions were defined accordingly. ‘Retained’ interactions are lost interactions that are higher in differentiated cohesin-depleted (IAA+RA-treated) cells than in normal (RA-treated) cells using the same criteria. Other lost interactions were categorized as ‘resolved’. Each bait that was involved in at least one retained interaction was defined as ‘Retained’, whereas baits that were involved in at least one resolved interaction but in no retained interaction, were defined as ‘Resolved’.

#### Definition of retained and resolved interaction anchors

For Figures 4 and 5, retained and resolved interaction anchors were defined as restriction fragments on both sides of a retained or a resolved interaction (i.e. both bait and otherEnd). For transcription analysis in Fig. 5, all anchors that overlapped both resolved and retained interactions, were assigned ‘retained anchors’. For footprinting analysis in Fig. 4, H3K27me3 peaks within 1kb of retained and resolved interaction anchors were defined as retained or resolved interaction anchors, respectively.

#### Analysis of cohesin-dependent rewiring

To ensure robust analysis and avoid an overestimation of the number of acquired interactions, fragment-level interactions were first merged if they were within two restriction fragments of each other using the PostChicago function aggregatePeaks() with the argument dis=2. This distance was chosen empirically upon inspection of Capture-C profiles. Aggregated interactions were then overlapped with fragment-level interactions and assigned either ‘gained’ or ‘lost’ if they overlapped at least one gained or lost fragment-level interaction. Interactions that overlapped both types of interactions were assigned ‘unclear’. All other interactions were defined as ‘stable’.

#### Native calibrated ChIP-seq analysis

Native calibrated ChIP-seq analysis was performed as previously described^62^. Briefly, paired-end reads were aligned using Bowtie2 v2.3.5.158^63^ (with “–no-mixed” and “–no-discordant” options) against concatenated mouse and human genome (mm10 and hg19). PCR duplicates were filtered out using SAMtools v1.659^64^ and Sambamba v0.7.166^65^. Only paired reads were retained. Following initial visualization and clustering using Deeptools v3.5.167^67^ to ensure a high correlation (cor>=0.9) between replicates, reads were downsampled to smallest library size and re-assessed for correlation again.

For visualization on IGV68, replicate reads were downsampled and read pile-ups generated using MACS2 v2.1.2.169^68^.

Peak calling was performed using the macs2 callpeak function from MACS2 with native ChIP input samples as control and non-downsampled native ChIP reads as input and the option -- scale-to small and --broad. The following parameters were used: H3K27ac:-q 0.1, --broad-cutoff 0.1, peaks were merged at 1kb, H3K27me3:-q 0.1, --broad-cutoff 0.15, --max-gap 250. The peaks were called individually for each replicate. The peaks from different replicates were subsequently merged using the reduce() function from GenomicRanges v1.30.360 to generate peak intervals. Depending on the number of replicates in a sample, only those peak intervals were retained that were identified in at least 2/3 or 3/4 biological replicates. Final peak sets were obtained by combining these filtered peak intervals called for individual samples using reduce().

For quantitative analysis, fragments overlapping peaks were counted in non-downsampled bam files using the summarizeOverlaps() function from the R package GenomicAlignments v1.14.160 with the parameters mode=’Union’ and paired=TRUE. For H3K27ac ChIP, spike-in calibration was incorporated by normalising the raw mm10 read counts using DESeq2 size factors, which were calculated based on the read counts for the set of unique hg19 intervals, as previously described^76^. For H3K27me3 ChIP, normalization was performed without the spike-in control, as its inclusion introduced a batch effect. To verify the absence of global changes, samples were also normalized using spike-in controls and analyzed at the level of individual replicates. As no global differences were detected, all subsequent analyses were conducted without spike-in normalization. The resulting count tables were then processed by DESeq2 v1.18.162^70^. The mean normalized read counts across replicates for each sample were calculated, with these being extracted from the DESeq2 results tables.

#### RNA-seq analysis

DESeq2 table with analyzed data was obtained from a previously published study^38^. A threshold for RA-expressed transcripts (-1) was empirically determined using the bimodal distribution of log2-transformed FPKMs derived from mean normalized read counts across replicates. Transcripts with a TSS within 5kb of a retained or a resolved interaction anchor were taken forward upon exclusion of transcripts whose transcription start and end sites exactly overlapped another transcript’s location.

#### Statistical analysis

Unless indicated otherwise, all comparisons were performed using the non-parametric Wilcoxon rank-sum test. Multiple-testing correction was performed by a multiplication of p-values <0.05 with the number of performed comparisons. Analysis of (n)cChIP-seq data was performed using DESeq2 and adjusted p values were used to determine significant values. Empirical p-values were calculated by conducting permutation testing of interactions.

#### R packages

Bioconductor^77^ was used to install R packages.

## Supplemental Information

**Table S1.** Summary of footprinting analysis for interaction loss, related to Figure 1.

**Table S2.** Capture-C oligo sequences.

**Table S3.** List of retained and resolved interaction anchors.

**Table S4.** Footprinting results at retained and resolved anchors in ES and RA cells, related to Figure 4.

**Table S5.** Gene expression data of the genes within 5kb from retained or resolved anchors, related to Figure 5.

---

**Figure S1.**
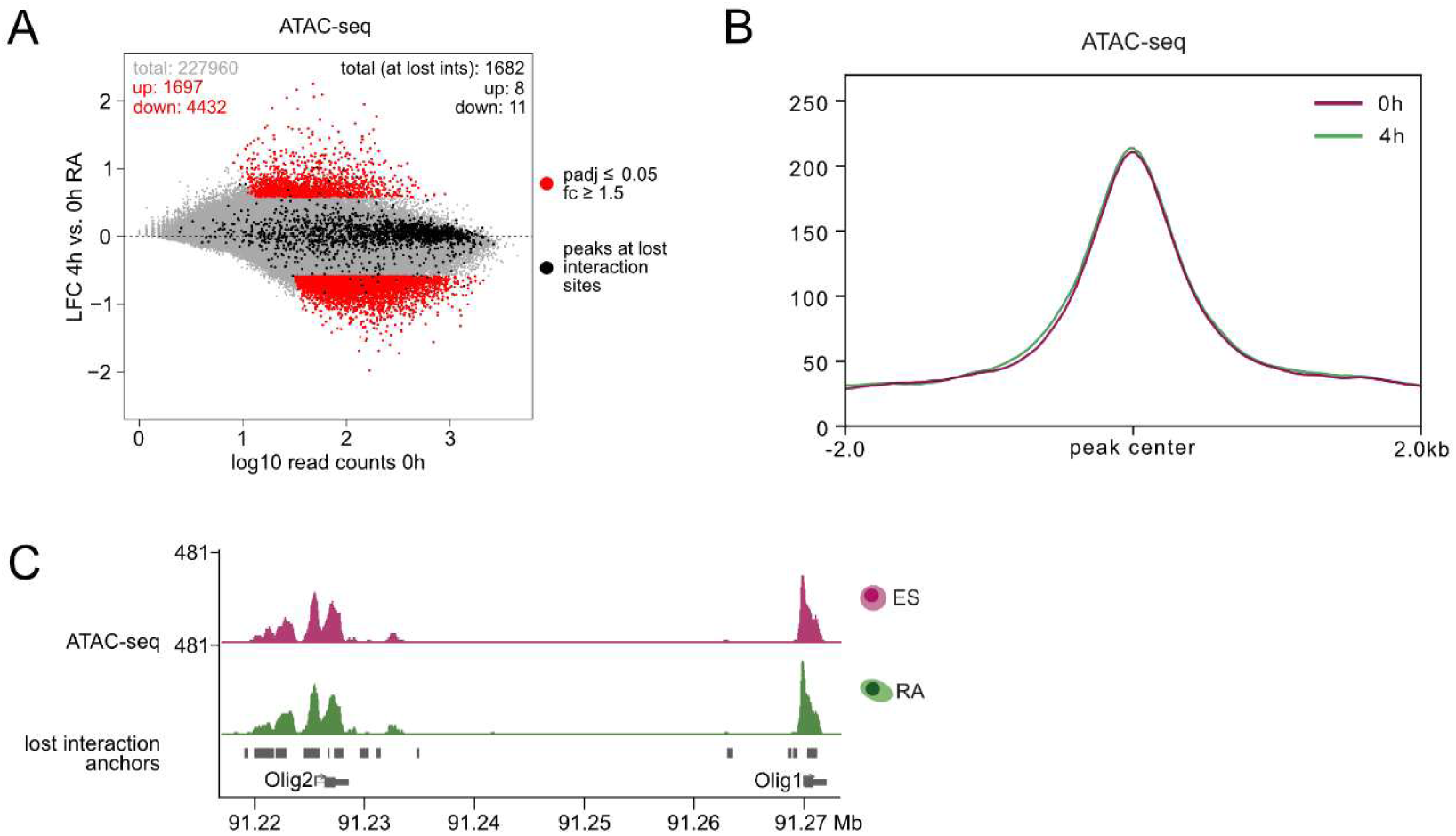
Unaltered accessibility at sites with lost interactions during early differentiation. (A) MA-plot showing the accessibility changes at ATAC peaks after 4h of RA induction (padj ≤ 0.05 and |fc| ≥ 1.5). **(B)** Aggregated ATAC peaks overlapping with genomic regions that lose interactions during the first 4h of RA induction (+/- 2 kb around the peak summit are shown). **(C)** Examples of representative ATAC peaks at 0h and 4h of differentiation overlapping with regions that lose interactions during that time.

**Figure S2.**
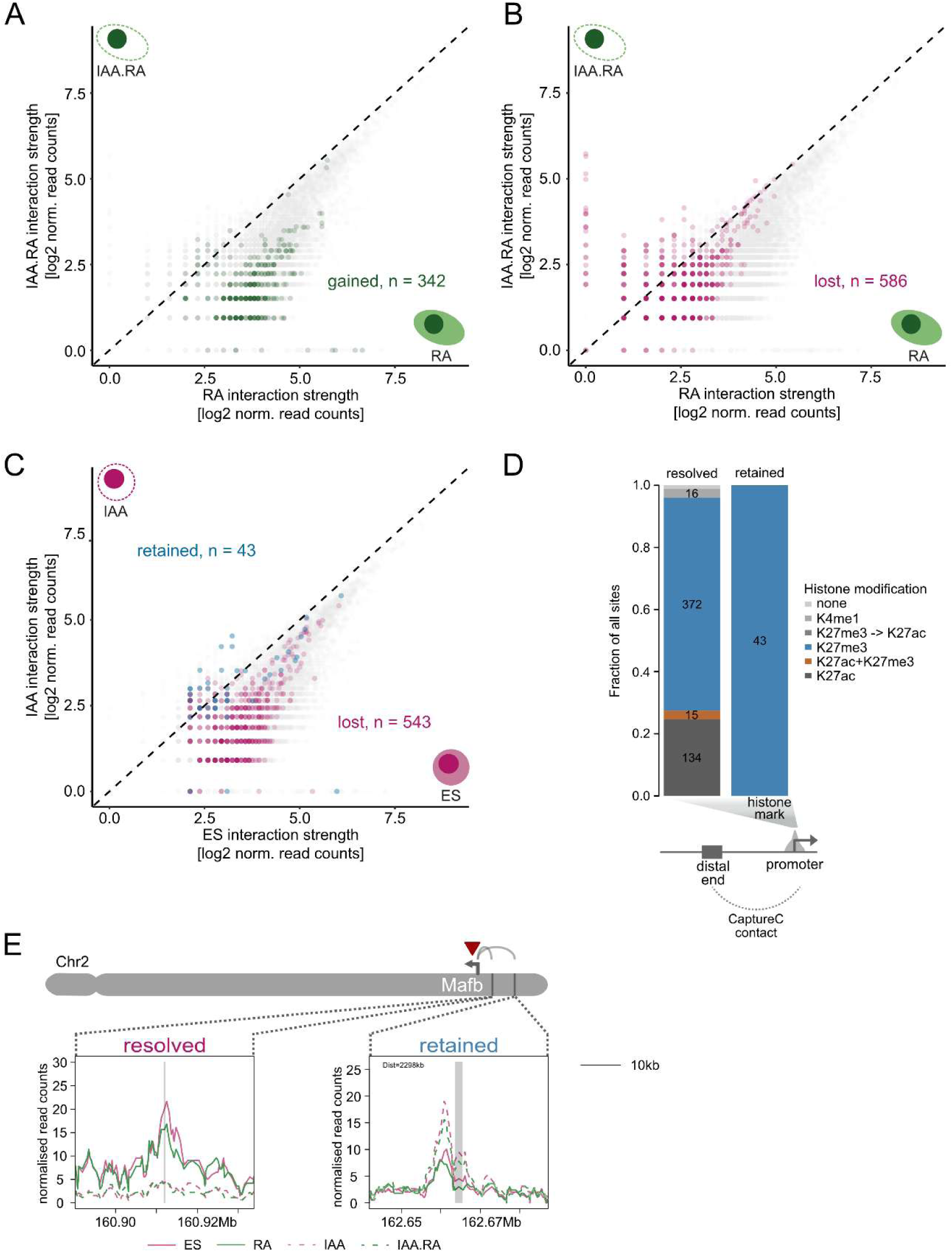
Cohesin is required for the efficient loss of a subset of promoter interactions during embryonic stem cell differentiation. (A-C) Scatterplots showing gained (A) and lost (B) promoter interactions in IAA.RA vs. RA, as well as interactions in cohesin-depleted vs wild-type ES cells (C). (D) Barplot depicting the enrichment of histone modifications^38^ at the promoters (baits) of resolved and retained interactions. (E) Example of a gene (Mafb) with both resolved (546kb away) and retained (2,298kb away) interactions.

**Figure S3.**
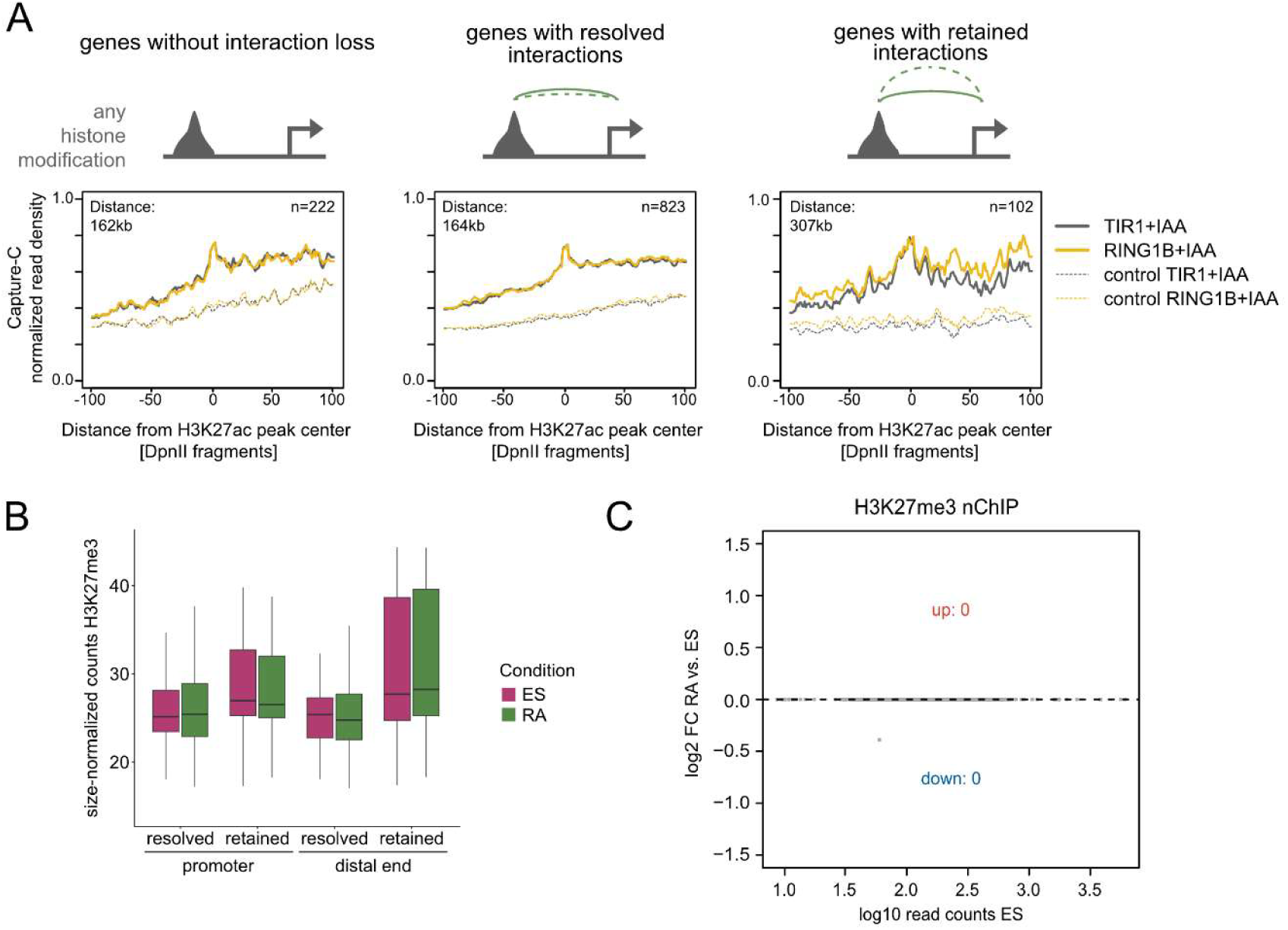
Retained interactions depend on Polycomb to the same extent as Polycomb-occupied resolved interactions. **(A)** Aggregated Capture-C plots demonstrating normalized interaction signal centred at H3K27ac peaks at resolved and retained sites before (grey) and after (yellow) RING1B depletion. Signal was normalized to the signal at the interaction summit of RING1B-AID. **(B)** Boxplots demonstrating peak size-normalized H3K72me3 levels at promoters and distal ends of resolved and retained interactions. **(C)** MA-plot comparison of H3K27me3 levels between RA and ES (DESeq2, |fold change| ≥ 1.5, padj < 0.05, n=4 replicates).

**Figure S4.**
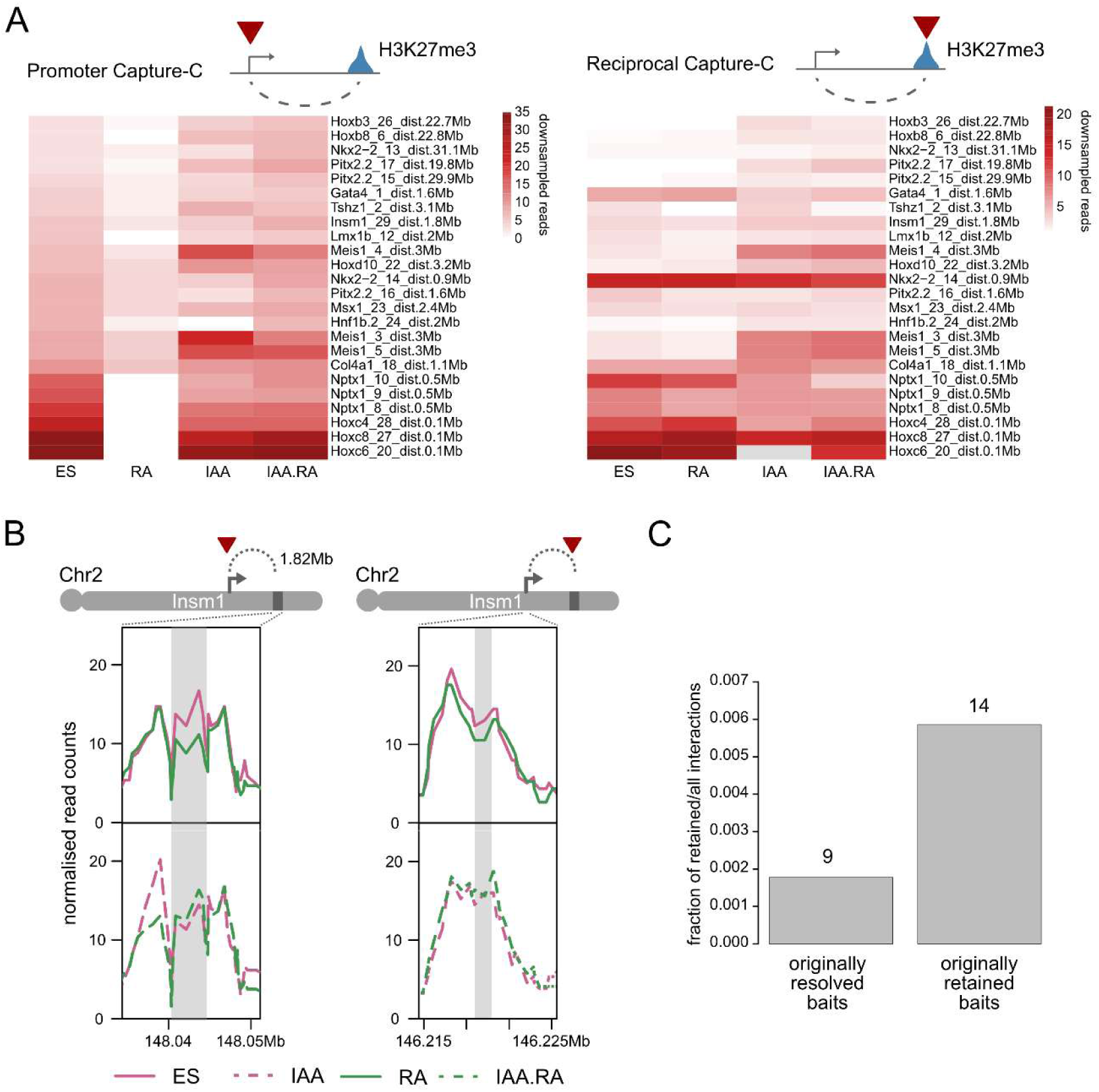
Reciprocal Capture-C validation. **(A)** Heatmaps depicting the interaction strength of retained sites in original promoter (left) and reciprocal distal (right) Capture-C experiments. **(B)** Example Capture-C signal is shown for the original promoter Capture-C of Insm1 (left) and reciprocal distal (right) Capture-C. Interactions are highlighted in grey. **(C)** Fractions of interactions assigned as retained based on the reciprocal Capture-C targeting sites that were previously involved in either resolved or retained interactions. Note that originally retained baits are enriched for retained interactions (p < 0.001), however additional retained interactions could now also be detected that involve sites with previously resolved interactions only.

**Figure S5.**
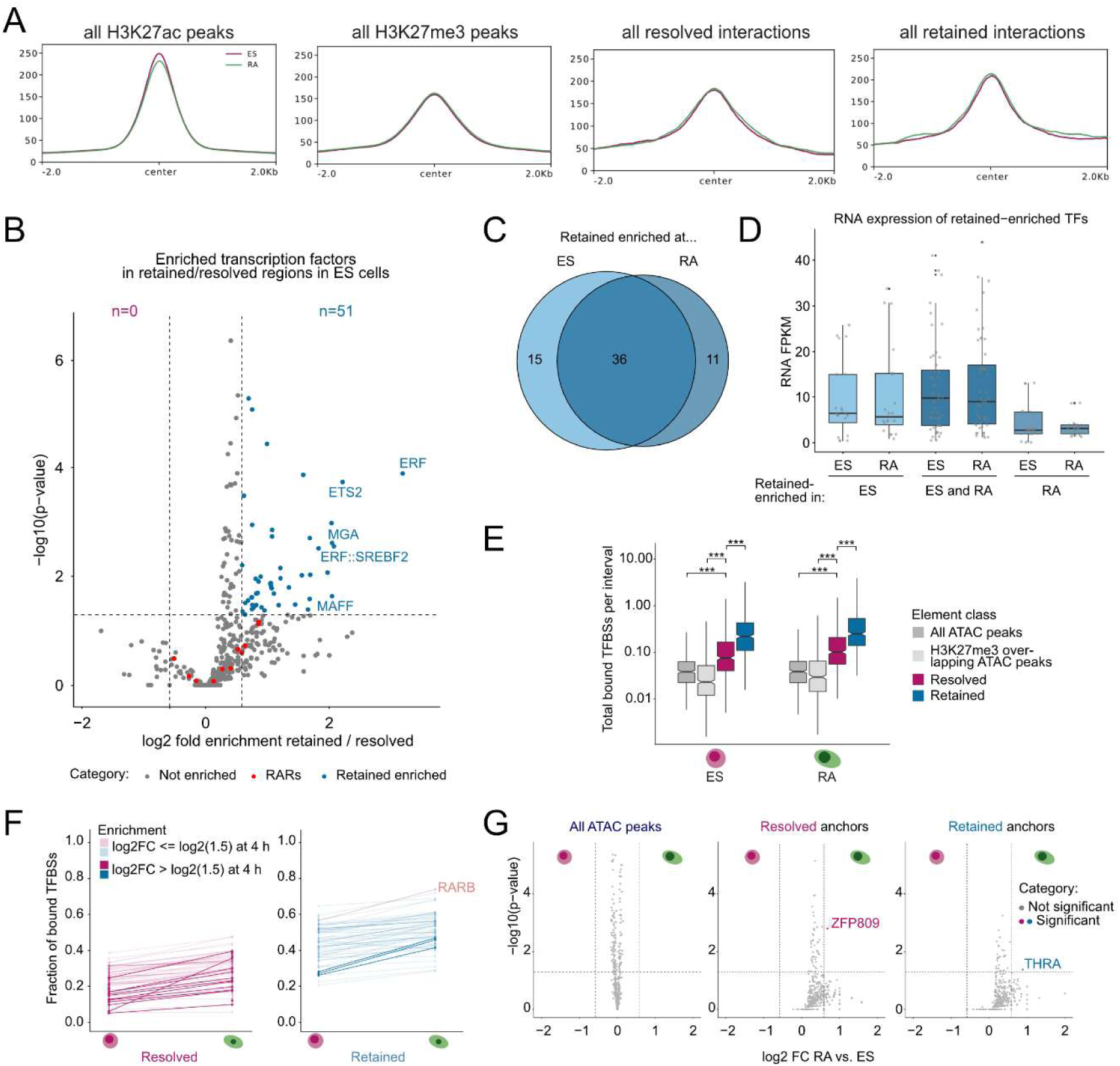
Retained interactions are enriched for sequence-specific transcription factor binding. **(A)** Metaprofiles of ATAC peaks in ES and RA overlapping H3K27ac peaks, all H3K27me3 peaks, and H3K27me3 peaks overlapping either resolved or retained elements (+/-2 kb around the peak summit) are shown. **(B)** Volcano plot demonstrating TF footprint enrichment in retained compared to resolved sites in ESCs. Top five TFs with the highest enrichment are labelled. **(C)** Venn diagram comparing the identity of TF footprints enriched at retained elements in ES and RA. **(D)** Boxplots show the expression of TFs enriched at retained elements in ES and RA. **(E)** Boxplots showing bound TFBSs per TF normalized by the number of intervals per subset (log scale). Adjusted p values are shown when significant: ∗p < 0.05, ∗∗p < 0.01, ∗∗∗p < 0.001. **(F)** Trajectory plots demonstrating the fractions of bound TFBSs at retained anchors in ES and RA. **(G)** Volcano plots demonstrating TF footprint enrichment in ES and RA at all ATAC peaks (left), resolved (middle) or retained anchors (right). Top five TFs with the highest enrichment are labelled.

**Figure S6.**
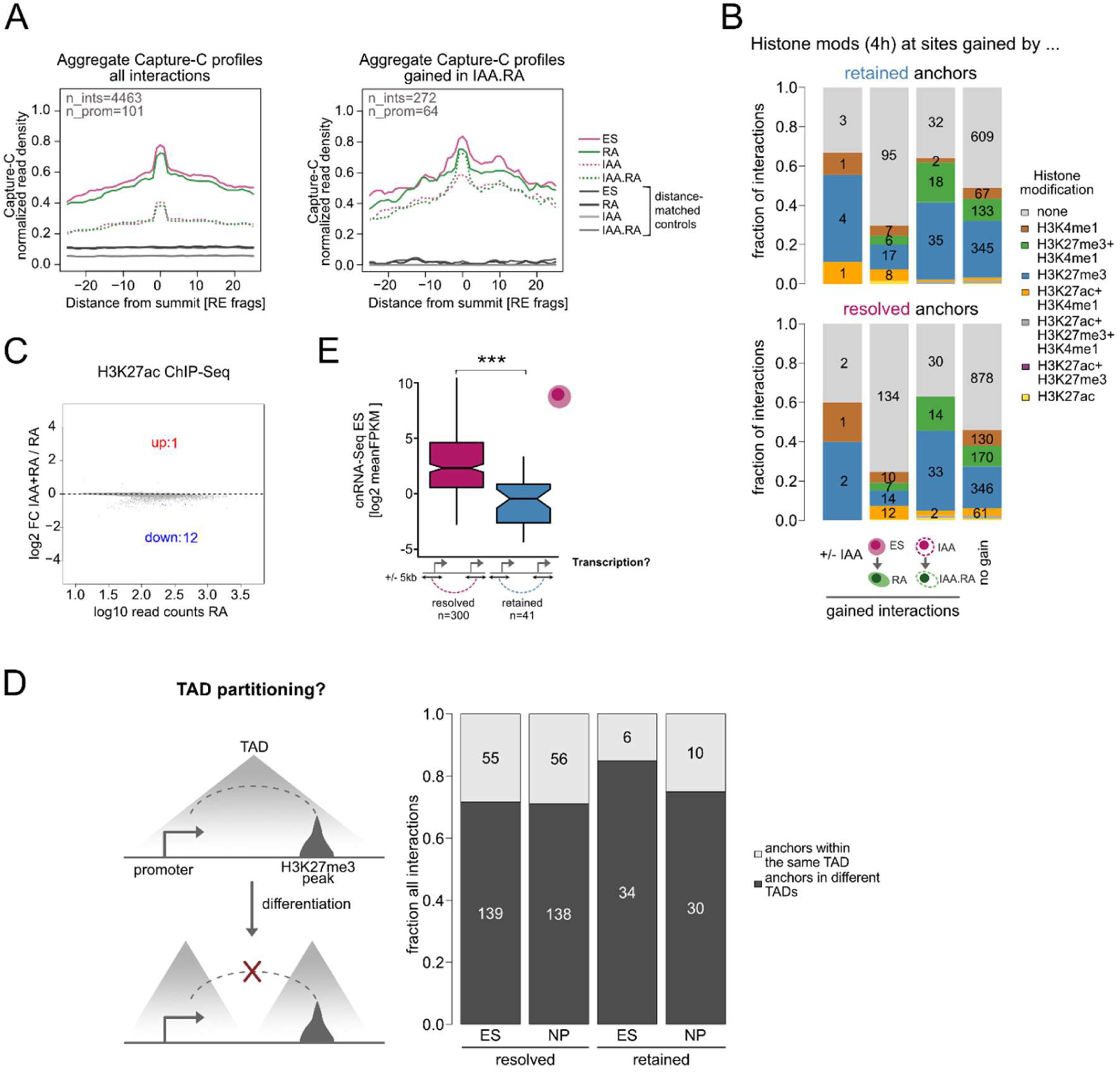
Retained interaction anchors undergo increased rewiring during the differentiation. **(A)** Aggregate Capture-C profiles of all detected interaction peaks (left) and those that are developmentally gained in cohesin-depleted cells (IAA-> IAA.RA, right). Signal is normalized to the signal at the interaction summit in ES. **(B)** Enrichment of histone modifications^38^ at the sites gained by either retained (top) or resolved (bottom) interaction anchors. **(C)** MA-plot comparison of H3K27ac levels between differentiated WT (RA) and IAA-treated (IAA.RA) cells (DESeq2, |fold change| ≥ 1.5, padj < 0.05, n=3 replicates). **(D)** Stacked barplots demonstrating the fraction of interaction anchors found in different TADs before (ES) and after *in vivo* differentiation (NP). TAD boundaries were downloaded from Bonev et al.^40^. Light grey: both anchors are located within the same TAD. Dark grey: both anchors are within different TADs. Note that TAD partitioning does not increase during the differentiation. **(E)** Boxplots showing ES transcription levels of genes expressed in RA that are located within 5kb of either resolved (red) or retained (blue) interaction anchors based on cnRNA-seq^38^. Adjusted p value: ∗∗∗p < 0.001.

